# The defining features of intrinsic transcription terminators

**DOI:** 10.64898/2026.05.29.728758

**Authors:** Robert A. Battaglia, Ebru Ermiş, Expery O. Omollo, Gene-Wei Li

**Affiliations:** Department of Biology, Massachusetts Institute of Technology; Cambridge, MA 02142, USA; Howard Hughes Medical Institute; Cambridge, MA 02142, USA

## Abstract

Transcription terminators are universal landmarks that delimit RNAs and tune downstream gene expression, yet the sequence rules that define them remain elusive. The classical model of bacterial intrinsic termination—an RNA hairpin followed by a uracil-rich tract—is incomplete, as stem-preserving mutations can abolish termination. By precisely mapping termination sites across 10^4^ sequences, we uncovered a previously unresolved feature required for intrinsic termination: dinucleotides positioned at both edges of the transcription bubble, resembling the elemental RNA polymerase pause signal. Together, hairpin, U-tract and bubble-edge sequences (HUB) account for most variation in termination efficiency and pinpoint bona fide terminators across diverse bacterial phyla. These findings establish HUB as the defining element of intrinsic terminators and provide a framework for decoding and engineering gene expression across genomes.

## Main Text

Factor-independent transcription termination is guided by U-rich sequences across all domains of life (*1–4*). In bacterial intrinsic terminators, a 7-8 nt U-rich sequence (U-tract) is preceded by a GC-rich hairpin. RNA polymerase (RNAP) pauses when the U-tract is within the RNA-DNA hybrid of the transcription bubble, facilitating hairpin folding into the hybrid, ultimately leading to transcript release and bubble collapse (*5–8*). Surprisingly, these hallmark features of intrinsic terminators are not sufficient to terminate transcription: mutations in the hairpin stem that preserve base-pairing complementarity can demolish terminator activity (*9*). Furthermore, models incorporating the hairpin and U-tract explain less than half of the variation in termination efficiency (TE, the fraction of times RNAP is successfully terminated by a given sequence) (*10*, *11*). Therefore, important features of intrinsic termination remain uncharacterized.

Intrinsic terminators are not merely endpoints of transcription, but pervasive regulatory elements that shape gene expression (*12*). Bacterial operons are frequently punctuated by intergenic terminators whose readthrough controls the expression stoichiometry among neighboring genes (*13*, *14*). Many terminators are also coupled to riboswitches and attenuators to mediate environmental or metabolic responses (*15–17*). Because intrinsic terminators are ubiquitous gene-expression landmarks in all domains of life, decoding their rules will shed light on broad principles of genome-wide regulation.

Uncovering necessary features required by intrinsic terminators is complicated by the repetitiveness of the U-tract, which confounds alignment in the absence of a point of reference. The exact location where RNAP terminates ultimately determines this point—each potential termination site corresponds to a different hairpin, RNA-DNA hybrid sequence, and positioning of the U-tract within the 8-11 bp hybrid (Fig. 1A) (*18–20*). Although termination sites and their TEs can be measured one-sequence-at-a-time using gel-based assays (*21*, *22*), it is difficult to systematically examine many different sequences to reveal general principles beyond amorphous hairpins and U-tracts. We reasoned that phasing each sequence relative to its termination site and treating each site as a distinct terminator would give us the resolution to uncover hidden features driving intrinsic termination.

**Fig. 1.**
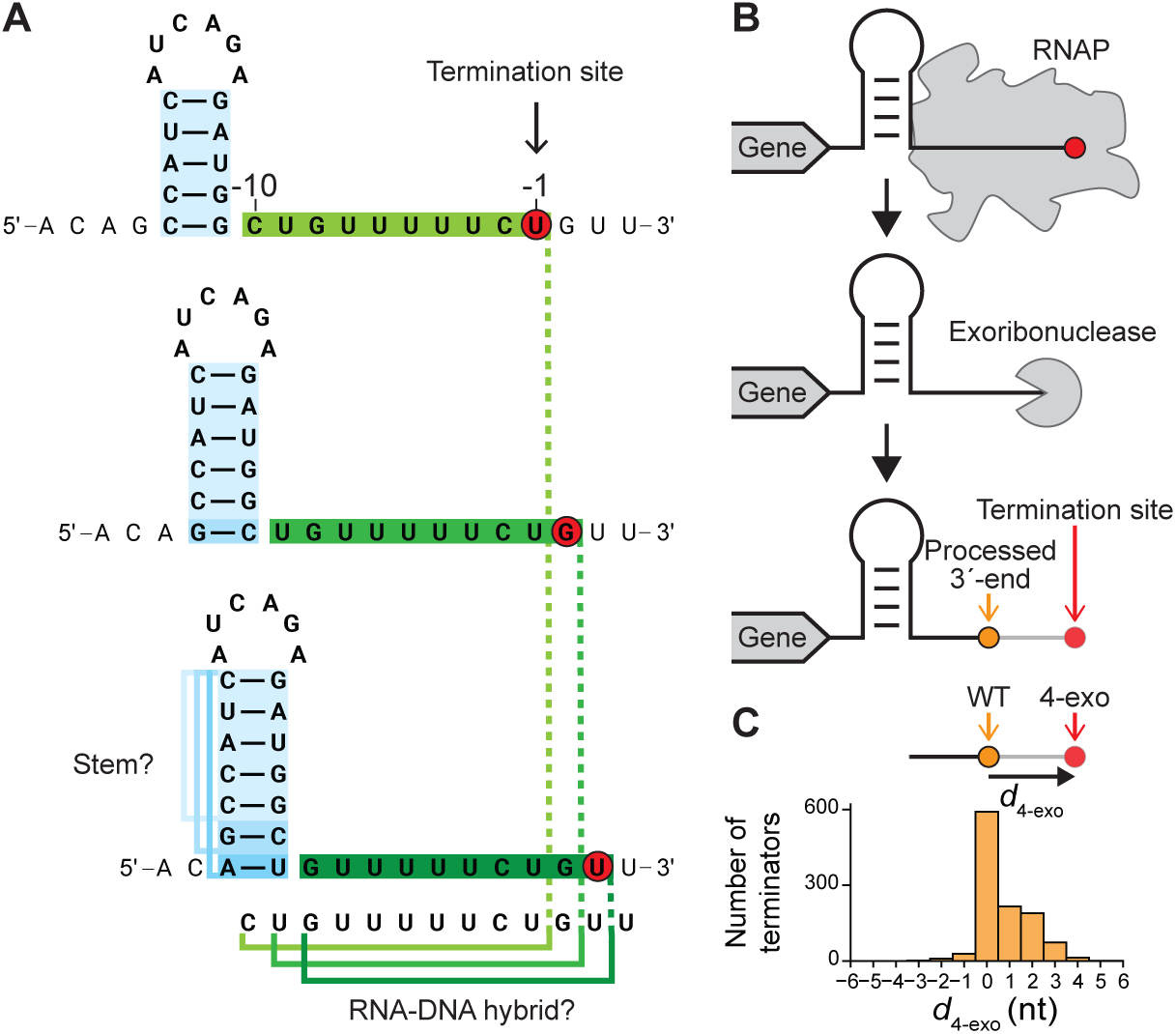
Identifying termination sites is important for resolving features of intrinsic terminators. (**A**) Assigning terminator features is challenging without knowing the termination site. Different termination sites (red circle) correspond to different RNA hairpin stems (blue) and regions in the RNA-DNA hybrid (green). The sequence is from the *B. subtilis yrhC* terminator. (**B**) *In vivo*, termination sites (red) can be processed by 3′-5′ exoribonucleases, shifting measured 3′-ends (orange) upstream. (**C**) Histogram of nucleotide distances between 3′-ends measured in *B. subtilis* with (WT) and without (4-exo) four major exoribonucleases (*d*_4-exo_).

### High-throughput mapping of termination sites

Although many terminators across the transcriptome have been mapped using 3′-end sequencing (*13*, *14*, *23–28*), we reasoned that the exact point of termination is likely different from the position of 3′-ends *in vivo*. One of several ways RNA 3′-ends can be altered *in vivo* is 3′-5′ degradation by endogenous exoribonucleases, which can proceed for several nucleotides until hindered by a hairpin structure (Fig. 1B) (*13*, *29–32*). Indeed, when we examined terminator-adjacent transcript boundaries from a *Bacillus subtilis* strain (4-exo) lacking most 3′-5′ exoribonucleases (PNPase, RNase R, RNase PH, and YhaM) (*28*, *33*, *34*), nearly half (44%) of the 3′-end positions are shifted downstream from the wildtype positions by at least one nucleotide (Fig. 1C). Therefore, 3′-ends measured *in vivo* are not adequate for elucidating the sequence contribution to TE.

To query the point of termination for thousands of intrinsic terminators without interference from post-transcriptional processing, we combined 3′-end sequencing with a reconstituted *in vitro* transcription reaction using a large pool of DNA templates. In the reaction template, each terminator and its surrounding sequence (60 nts upstream and 30 nts downstream) are inserted between a promoter and an invariant strong terminator downstream (Fig. 2A). After transcription, the 3′-ends of released RNAs are ligated to an adaptor and converted into a DNA library for deep sequencing (*35*). The likelihood of terminating at each position in each template can be assigned by counting corresponding 3′-ends (Fig. 2BC). Applying this method to the classic λtR2 terminator recapitulates the two prominent sites of termination, showing a weaker signal at the first site (historically labeled as U7) and a stronger signal at the second site (U8) (Fig. 2D) (*21*, *36*).

**Fig. 2.**
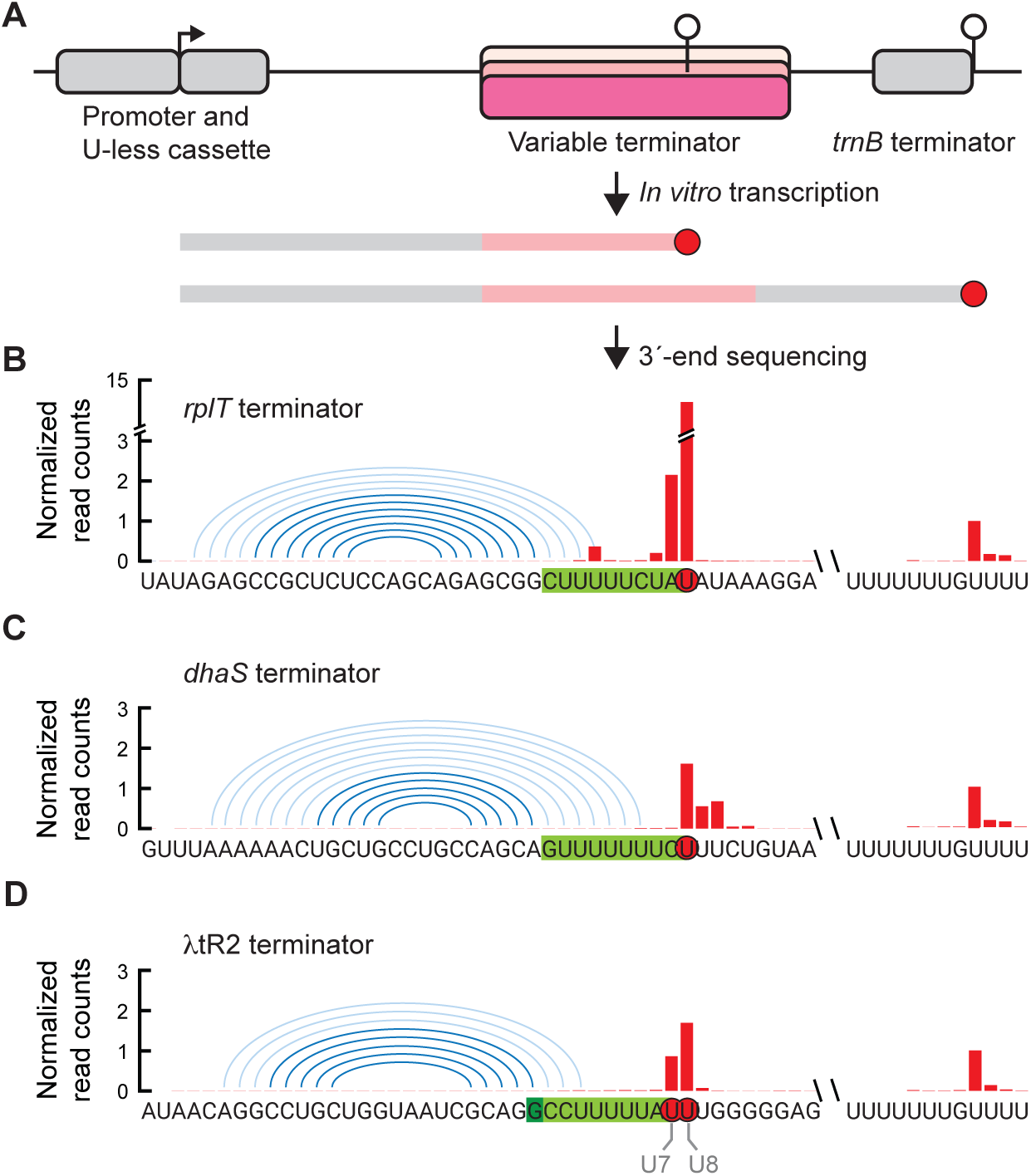
Pooled *in vitro* assay maps sites of termination. (**A**) Design of template libraries and sequencing approach. Each template includes a promoter and a downstream sequence without uridines (U-less cassette) that enables single-round transcription. Different terminator sequences are inserted between the constant promoter and downstream terminator (*trnB*). After *in vitro* transcription, the 3′-ends of RNAs (red) are ligated to an adapter and sequenced. (**B**-**D**) 3′-end sequencing results for strong (**B**, *rplT*) and weak (**C**, *dhaS*) *B. subtilis* terminators transcribed by *B. subtilis* RNAP and the canonical λtR2 phage terminator transcribed by *E. coli* RNAP (**D**). Read counts are normalized to the number of reads mapped to the strongest termination site at the downstream *trnB* terminator. RNA-DNA hybrid sequences corresponding to dominant termination sites are highlighted in green. Hairpin base pairs are labeled by dark blue arcs, whereas potential base pairs that require RNA-DNA hybrid melting are shown as light blue arcs. The distribution of 3′-end sequencing reads across the two termination sites of the λtR2 terminator (U7 and U8) are consistent with gel-based methods (*36*).

Using this approach, we examined the activity of RNAP across 2,008 intrinsic terminators from *B. subtilis* and *E. coli*. Only 24% of *B. subtilis* terminators transcribed by the *B. subtilis* RNAP generate identical 3′-ends *in vivo* and *in vitro* (Fig. S1AB). For most (72%), the prominent points of termination *in vitro* are downstream of the 3′-ends observed in the wildtype (Fig. S1BC). *In vitro* termination sites are more similar to sites in the 4-exo, but a substantial fraction (45%) is shifted further downstream, suggesting that additional processes can shape transcript boundaries *in vivo* (Fig. S1CD) (*28*, *37*). Consistent with established mechanisms of termination, *in vitro* termination sites correspond to more stable RNA hairpins and less stable RNA-DNA hybrids in the RNAP elongation complex than those defined by the *in vivo* 3′-ends (Fig. S1EF). Taken together, these results suggest our high-throughput approach reports authentic points of termination.

### Resolving site-specific termination efficiency

Many terminators generate 3′-ends at multiple adjacent positions, indicating that termination takes place at consecutive sites with distinct TEs (*yerC,* Fig. 3A). We found that nearly half of the terminators (41%) have at least 2 sites that contribute to >20% of the overall termination activity (Fig. 3B). As noted above, multi-site termination has been observed for the λtR2 terminator (*21*) and is consistent with the fact that successive sites in any given terminator sequence can have corresponding hairpin and U-tract features (Fig. 3A). In fact, the strong λtR2 terminator should be viewed as a composite of two moderate terminators, instead of a single strong terminator (e.g. *rplT* in Fig. 2B).

**Fig. 3.**
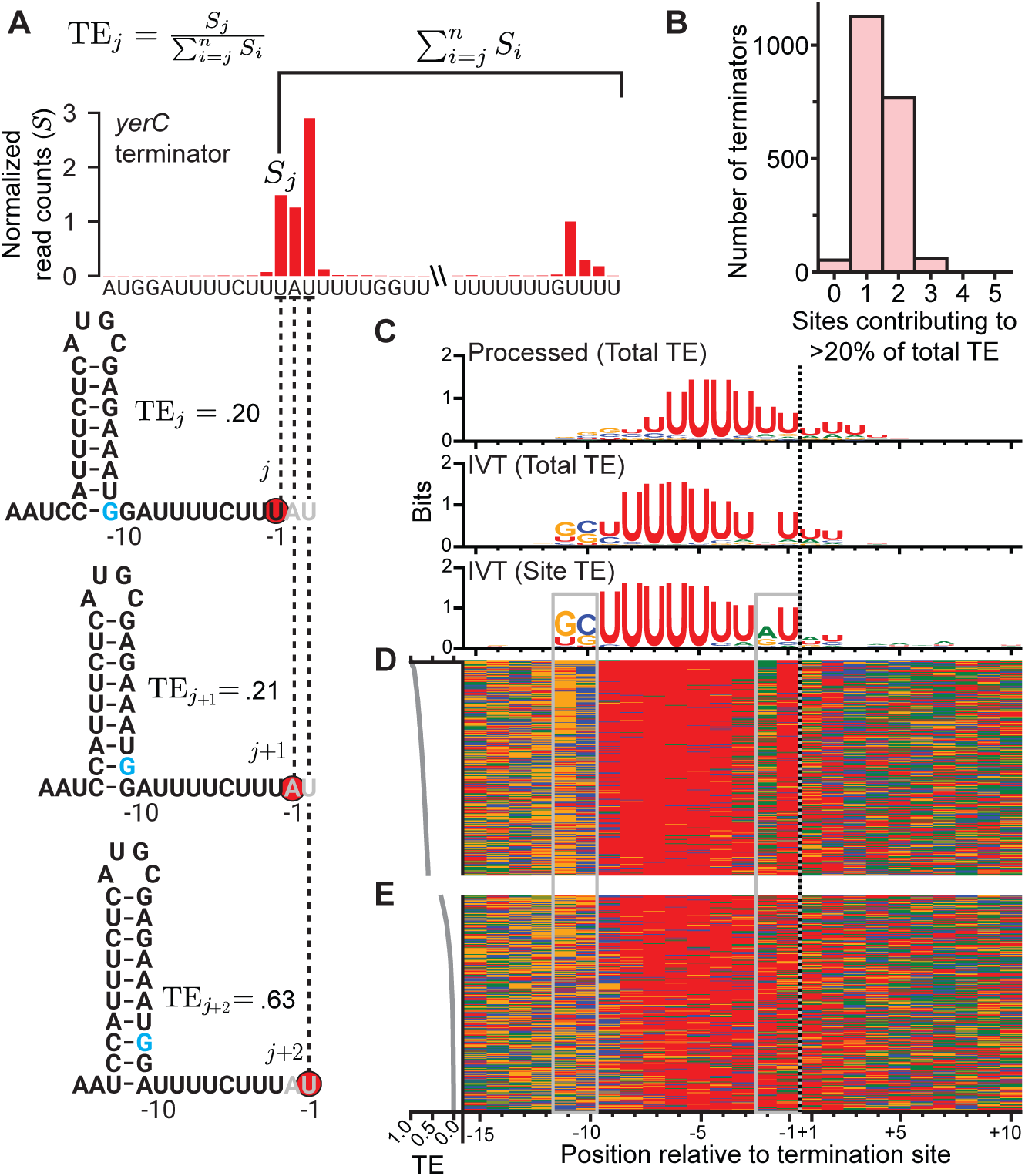
Quantifying site-specific TE reveals common bubble-edge motifs. (**A**) The total TE of a given terminator is a composite of the site-specific TEs of distinct terminator sequences associated with each termination site. The *yerC* terminator has multiple termination sites (red circles), each with unique hairpin and hybrid sequences. -11 position for the most upstream termination site colored blue. The site-specific TE for each termination site (TE*_j_*) is calculated as the ratio of reads at site *j* to sum of reads at site *j* and all downstream sites (inset). (**B**) Histogram of terminators with different numbers of sites contributing to total TE. The number of sites that contribute to at least 20% of total TE is counted for each terminator. (**C**) Sequence logos for strong terminators (TE > 0.9) generated relative to *in vivo* (processed) or *in vitro* (IVT) 3′-ends (designated position -1). Strong terminators are determined using total TE or site-specific (Site) TE (label in parentheses). Dotted line indicates the 3′-end position. (**D**) 300 representative sequences of the strongest termination sites (TEs ranging from 1.0 to 0.6) recapitulating the GC(U)_7_RU motif found in (C). Color-coded sequences -15 – +10 relative to termination site (-1) are arranged in descending order by TE. Each row corresponds to a termination site sequence with the color in each column representing the base identity at each position (green: A, blue: C, orange: G, red: U). Site TE across rows is plotted on the left. Grey boxes indicate locations of - 11/-10 and -2/-1 nucleotides. (**E**) 300 representative sequences of the weakest termination sites with TE ranging from 0.3 to 0.01.

To match the correctly phased sequence with TE at each site of termination, we quantified site-specific TE as the fraction of RNAPs that terminated at a site *j* (given by 3′-end signal, *S_j_*) out of all RNAPs that reached site *j*. The number of RNAPs reaching site *j* was estimated as the sum of 3′-end signals at and downstream of that position (∑*^n^_i=j_ S_i_*), since every RNAP that reaches *j* must terminate either at *j* or further downstream (Fig. 3A) (*21*). As a proof-of-principle, we used our approach to examine the activity of *E. coli* RNAP on the λtR2 terminator (*21*). We observed that the relative TE at the two prominent termination sites was quantitatively consistent with previous gel-based measurements across an array of mutations (Fig. S1G) (*36*). Site-specific TE is sensitive to nucleotide concentrations, the presence of NusA, and to a much lesser extent, NusG (Fig. S2ABC) (*14*, *21*, *23*, *38*, *39*). However, the rank order among the ∼13,000 – 17,000 TEs remains largely consistent across conditions (Fig. S2DEF). This large, spatially resolved dataset of site-specific TEs provides an opportunity to reveal how precise sequence features at the RNAP-nucleic acid interface affect the fate of elongation.

### GC(U)_7_RU motif enriched at strong termination sites

Properly phased nucleotide sequences at the strongest termination sites revealed a consensus sequence that augments the current definition of intrinsic terminators (Fig. 3C). A signature for the canonical 7-nt U-tract starts at the -9 position relative to the RNA 3′-end (-1), not the -8 starting point inferred by sequence-gazing without phasing (Fig. 3C and (*20*, *40*)). Importantly, we identified two additional features flanking the (U)_7_ region that are enriched among sequences of the strongest sites: a GC dinucleotide starting at the -11 position and a RU dinucleotide (AU and GU) starting at -2 (Fig. 3CDE). These features reside at the edges of the transcription bubble, and emerge only after removing the confounding *in vivo* 3′-end processing and resolving site-specific TE (Fig. 3C). The overall GC(U)_7_RU motif also emerged when we used *E. coli* RNAP in our *in vitro* assay (Fig. S2G), suggesting this motif is a broadly conserved signature of bacterial terminators.

Intriguingly, the GC(U)_7_RU motif resembles a U-tract sandwiched by the bipartite consensus of elemental RNAP pause sites (GS(N)_7_RY, where S stands for C and G, and Y for U and C) (*41*, *42*). An elemental pause is a transient off-pathway state common to elongating RNAPs in many organisms (*43*). RNA-DNA pairing at the -11/-10 positions in particular has been implicated in slowing forward translocation of RNAP (*44–46*). For intrinsic terminators, a U-tract pause is thought to be the first step towards transcription termination, providing time for hairpin formation and the subsequent steps towards RNA dissociation (*5–7*, *47–50*). The observed U-tract pause could be further mediated by elemental pause-like signals, which are an essential element of strong terminators because of the crucial kinetic window pausing provides for entering the termination pathway.

### Bubble-edge sequences are an integral feature of intrinsic termination

To determine whether the GC and RU motifs at the edge of the transcription bubble are necessary for transcription termination, we designed mutational libraries for several intrinsic terminators to examine the effect of mutating the regions surrounding the U-tract (Fig. 4A). In these libraries, each variant is associated with a unique upstream barcode so that terminated products from different DNA mutants can be distinguished in our 3′-end sequencing approach (Fig. 4B).

**Fig. 4.**
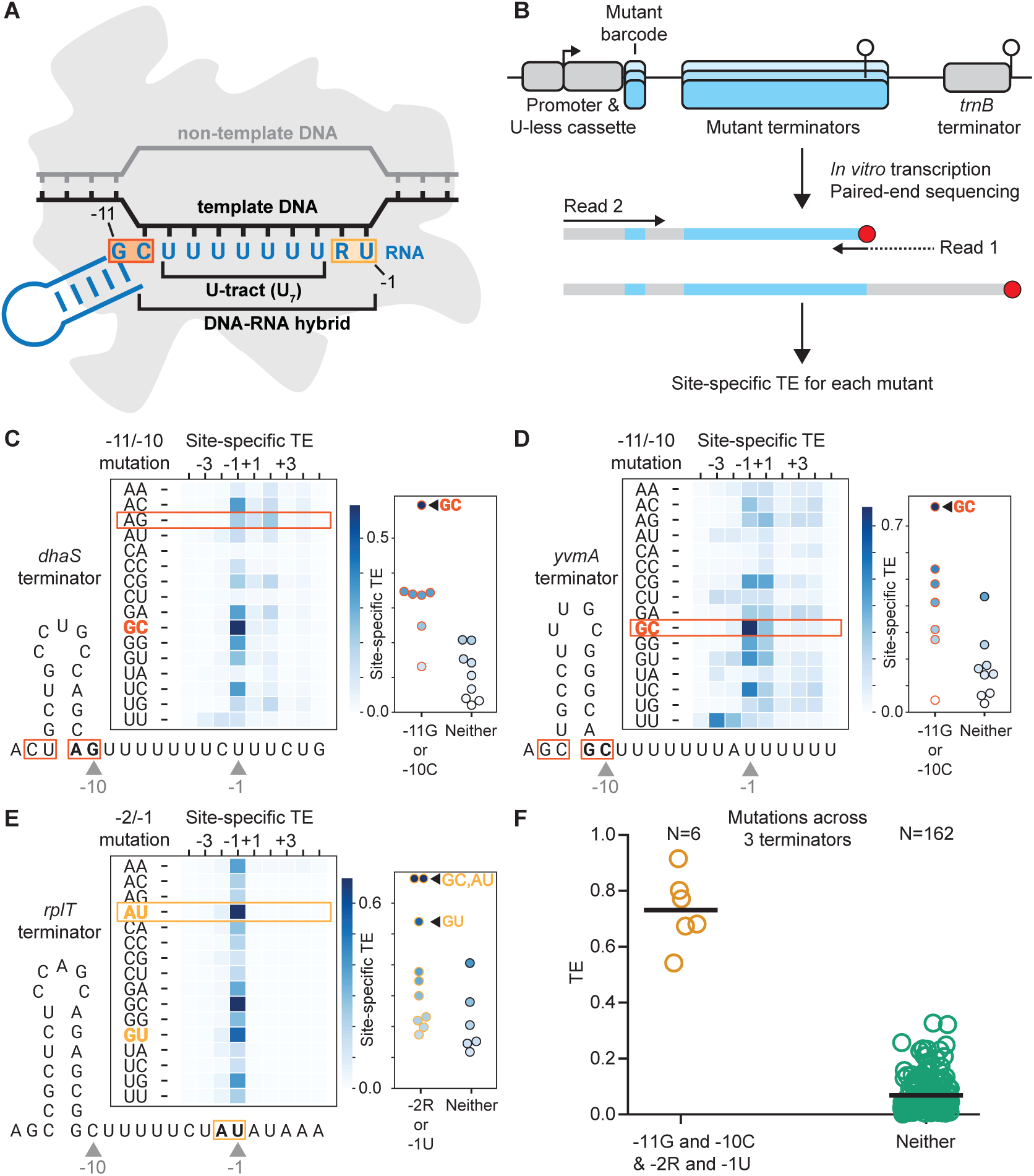
Mutagenesis of the bubble-edge motif demonstrates that it is necessary for terminator function. (**A**) The GC(U)_7_RU motif places the -11/-10 GC (red box) and -2/-1 RU (orange box) dinucleotides on the edges of the transcription bubble. The termination site is position -1. (**B**) Barcoded 3′-end sequencing distinguishes terminated and read-through products from different mutant terminators. (**C**-**E**) Comprehensive mutagenesis of the -11/-10 and -2/-1 positions to evaluate the necessity of the bubble-edge sequence motifs for termination. Mutations were introduced to a native *B. subtilis* terminator lacking GC at -11/-10 (*dhaS*, **C**), a terminator with GC at -11/-10 (*yvmA*, **D**), and a terminator with RU at -2/-1 (*rplT*, **E**). Mutagenized sites are indicated by red (-11/-10 and the complementary mutations in the upstream stem) or orange (-2/-1) boxes on the sequence diagram. WT residues are boxed on heatmaps, whereas consensus residues are colored and bolded. Heatmaps display site-specific TEs in a 9-nt window around the -1 site for each mutant. Right inset: dot plots of site-specific TEs at -1 position for mutants with or without any motif matching sequences. (**F**) Comparison of mutants with and without the bubble-edge sequence motif. Dot plot comparing TEs for *dhaS*, *yvmA*, and *rplT* mutants that perfectly match the -11G/-10C/-2R/-1U motif (-11G and -10C & -2R and -1U, yellow) to mutants that have no match to the motif (Neither, green).

We first focused on the GC motif at positions -11/-10. Because both positions base pair with upstream sequences in the terminator hairpin (Fig. 4A), all mutations at -11/-10 were complemented by mutations in the upstream stem to preserve hairpin structure. Starting with a weak termination site (*dhaS*: TE=21%) that has AG at the corresponding -11/-10, mutation to the consensus GC causes the greatest increase in TE across all possible substitutions (TE=59%) (Fig. 4C). Conversely, starting with a strong termination site with GC at -11/-10 (*yvmA*: TE=77% and *rplT*: TE=72%), any mutation at these positions decreases TE (Fig. 4D and Fig. S3B, respectively). Gel-based TE measurements of *rplT* mutants are consistent with these findings (Fig. S3A). Importantly, mutants without -11G or -10C—even though a strong hairpin and U-tract are maintained—barely terminate at the original position (median TE=6%) (Fig. 4D). Combined with mutational analyses across 15 additional terminators (Supplementary Text, Fig. S5), these results establish the GC motif at the interface of the hairpin and RNA-DNA hybrid as a key requirement for intrinsic terminators.

We next verified the contribution of the RU motif at the -2/-1 positions. In agreement with our sequence logo, mutations away from a U at the -1 position generally reduced TE across the terminators we assayed (*dhaS*, *yvmA*, and *rplT*, Fig 4E, S3CD). Likewise, the combination of a -1U with a purine (A or G) at -2 is associated with the strongest TE. Further supporting the role of RU, we observe that mutations resulting in an RU motif at -1/+1 (*dhaS* and *yvmA*, Fig S3CD) increase TE at the +1 site. We note that GC at -2/-1 in the *rplT* terminator has a TE value similar to AU (TE=68%) and greater than GU (TE=54%). This result is consistent with our consensus sequence, which shows a small but noticeable information content for C at the -1 position. Overall, mutations with the highest TE values are strongly enriched for -2R or -1U across the 18 terminators we tested (Fig. S5). Together, these results indicate that a U-tract interrupted by a purine at -2 substantially enhances termination.

To further test the necessity of GC and RU motifs, we expanded the mutational analysis by generating all possible combinations of mutations at these four positions (and the matching positions upstream to preserve hairpins) across 3 terminators (*dhaS*, *yvmA*, and *rplT*, 768 mutants). Mutants that contain none of the preferred nucleotides at -11/-10 and -2/-1 (N=162) have negligible TE (median TE=4.3%, Fig. 4F), whereas mutants with a perfect match to the motif strongly terminate (N=6, median TE=73%). Therefore, hairpin and U-tract structure alone are insufficient for intrinsic termination—a defining sequence feature at the edges of the transcription bubble is also necessary.

The necessity of the GC(U)_7_RU motif also explains a longstanding mystery about the extensively characterized λtR2 terminator. Early work showed that mutations preserving complementarity in the λtR2 stem are surprisingly detrimental to termination, despite having minimal effects on hairpin stability (*9*). The λtR2 stem-loop contains a GC dinucleotide at positions -11/-10 (relative to its most prominent site of termination, U8). Based on our findings above, we expected this dinucleotide to be critical for termination. Indeed, in our stem-preserving λtR2 mutation library, transcribed using either *E. coli* or *B. subtilis* RNAP, we found diminished termination activity when the -10C is disrupted, especially when combined with disruption of -11G (Fig. S4AB). The effects of other mutations to λtR2, including changes in TE and shifts in termination site, can also be explained by our model (Supplementary Text, Fig. S4). Altogether, these results establish that intrinsic terminators should be identified by a tripartite feature—<u>h</u>airpin, <u>U</u>-tract and <u>b</u>ubble-edge sequences (HUB).

### An integrated model for intrinsic terminators across bacteria

We next evaluated the prevalence of HUB-based termination across diverse bacteria. To do so, we first developed machine-learning models for predicting TE from sequences and then examined whether inclusion of bubble-edge features improves terminator identification for distantly related genomes. For TE prediction, even in *E. coli*, it has remained difficult to distinguish strong terminator sequences from weak or non-terminating ones using the classic definition of intrinsic terminators (hairpin and U-tract, herein referred to as HU) (*10*, *11*). We incorporated the HUB features and our large dataset of site-specific TE measurements to train TE-prediction models. For each of the 22,369 termination sites across endogenous terminators, we parametrized phased sequence features including both the HU features, as well as the newly identified bubble-edge motifs (B). Using models based on gradient boosting decision trees, we found that the classic features alone capture 57% of variations in TE (cross-validated R^2^, Fig 5A). This model’s predictions are slightly improved compared to previous TE prediction models (*10*, *11*), potentially enabled by our phased definition of hairpin and U-tract. In contrast to the HU-based model, a model that includes all three HUB features increases predictability to 68% (Fig. 5AB). Notably, 69% of predictions are within two-fold of the measured values (Fig. 5C). Furthermore, including sequence information from an expanded window (positions -41 to +19) only slightly increases predictability (R^2^=71%), suggesting that there is minimal remaining information beyond HUB. Similar results were found for RNAPs from both *B. subtilis* and *E. coli*, reactions with and without NusA, and at different nucleotide concentrations (Fig. S6). These results further confirm that the bubble-edge motifs are a key feature missing from our current definition of intrinsic terminators.

**Fig. 5.**
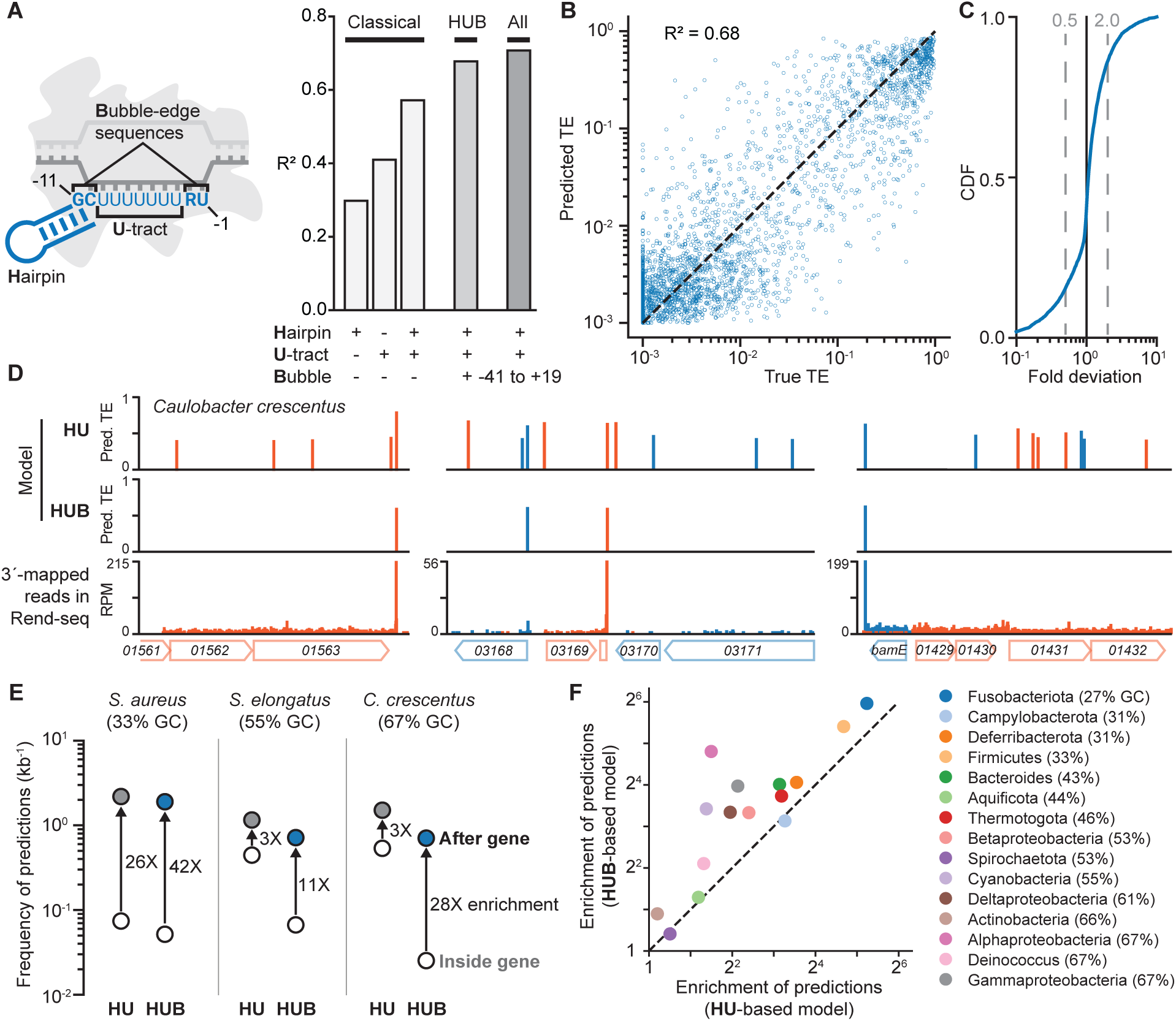
The generality of HUB (<u>H</u>airpin, <u>U</u>-tract, and <u>B</u>ubble-edge sequences) across bacteria. (**A**) Models that incorporate various HUB features for predicting TE. Left: schematic of HUB features and their positions relative to the termination site. Right: Bar graph of R^2^ values for statistical models trained with different feature sets. Classical feature set includes Hairpin and U-tract (Methods). The HUB feature set further includes Bubble sequence information (sequence of -11 to -1 region), whereas “All” features include sequence from an extended -41 to +19 region. (**B**) Predicted TEs are plotted against measured TEs (True TE). True TEs that are < 10^-3^ are set to 10^-3^. (**C**) Cumulative distribution function (CDF) of the fold-deviation of predicted TEs from measured TEs. 2-fold deviation indicated by grey dashed lines. (**D**) Representative genomic regions in *C. crescentus*, showing model TE predictions (HU or HUB), 3′-mapped reads in End-enriched RNA-seq (Rend-seq) (*13*), and gene annotations. Forward and reverse strand features are colored orange and blue, respectively. RPM: reads per million. Numbers inside genes are the unique identifiers following the prefix “CCNA_”, except for *bamE*. (**E**) Dot plots showing the frequency of terminator predictions in non-coding regions downstream of stop codons (after genes, filled circle) and inside genes (open circle). Three representative genomes with different GC content are shown. HUBfinder predictions are in blue and HUfinder predictions are in grey. (**F**) Scatter plots showing the enrichment of predictions after genes by HUBfinder or HUfinder across representative genomes from 15 different bacterial phyla (Table S3). Enrichment of predictions is calculated as the ratio of prediction frequencies after genes and inside genes. GC content of genome used for each phylum is in parentheses after phylum name in legend.

The TE-prediction models allow us to examine the generality of HUB across diverse bacterial genomes. If bubble-edge motifs are a key component of intrinsic terminators, a HUB-based model should be more accurate at identifying terminators than a HU-based one. Indeed, for the distantly related species *Caulobacter crescentus*, our machine-learning model that incorporates all HUB features (HUBfinder) successfully predicts terminators near previously mapped mRNA 3′-ends (Figs. 5D and S7) (*13*). Importantly, the predicted terminators are primarily located downstream of genes, in contrast to the numerous intragenic predictions made by a model based solely on HU (Fig. 5DE). The latter are likely false discoveries due to an incomplete feature set, a common challenge faced by previous terminator-calling algorithms (*51*, *52*). Across representative species from diverse phyla, we found strong enrichment of HUB-based predictions downstream of genes (Fig. 5EF). By contrast, HU-based predictions show substantially weaker enrichment, especially for species with GC-rich genomes where specifying terminators may rely on correctly placed -11G and -10C relative to the U-tract (Fig. 5EF). Notably, our model was trained with *B. subtilis* and *E. coli* terminator sequences and does not include data from the diverse organisms examined here. Therefore, these results illustrate the universality of HUB as the definition of intrinsic terminators in bacteria.

## Discussion

We have identified a third, crucial component of intrinsic terminator activity: the sequence of positions -11/-10 and -2/-1 at the edges of the transcription bubble. Why are these positions important for termination? The optimal motif surrounding the U-tract (GC(U_7_)RU) closely resembles the consensus elemental pause sequence (GS(N_7_)RY) (*41*, *42*, *53*). Pausing is thought to be the first step towards termination, providing time for the hairpin to fold in the RNAP exit channel (*1*). A few lines of evidence link elemental pausing and termination: (i) Mutations to the -2 and -1 positions that decrease termination in λtR2 have been shown to also decrease pausing (*6*). (ii) The half-translocated conformation that is a common feature of paused RNAPs from bacteria to mammals has also been observed in pre-terminated RNAP structures (*20*, *54–56*). The elemental pause sequence elicits pausing—and potentially a half-translocated state—in RNAPs from all domains of life (*41*), suggesting that stabilizing this state via template sequence could be a universal strategy for regulating transcription termination. Nevertheless, it is also possible that GC and RU at their respective locations have additional contributions to termination beyond affecting pausing. Further kinetic studies *in vitro* could help isolate the mechanistic roles of these sequence features and quantify their relative contributions to termination.

Although models that consider all HUB features are able to capture most of the variation in TE across terminators, unexplained differences remain. At certain terminators, the propensity for upstream sequences to fold into anti-terminator structures could also influence TE (*11*, *57*, *58*). Furthermore, our approach does not allow us to distinguish two proposed mechanisms of intrinsic termination: hypertranslocation and hybrid-shearing (*1*, *59*). In the context of hypertranslocation, the sequence downstream of the termination site could modulate TE by affecting how readily RNAP is moved forward (*40*, *60*). The contribution of these mechanisms could be investigated using an expanded mutational library together with differential presence of NusA and NusG, which are thought to favor distinct mechanisms of RNA release (*61*). Finally, further improvement in TE prediction could be achieved by augmenting the training data with larger-scale experimentation or evolutionary comparison (*62*).

Across bacterial genomes, transcription terminators are a fundamental building block for numerous regulatory elements, ranging from attenuators and riboswitches to programmed read-through systems. The discovery of HUB as the hallmark of intrinsic termination establishes a renewed framework for investigating these regulatory components. Furthermore, this discovery resolves a decades-old puzzle, explaining how hairpin-preserving mutations can nevertheless disrupt termination. More broadly, the ability to accurately identify terminators and predict their efficiency will transform our capacity to both decipher and engineer bacterial genomes.

## Acknowledgments

We thank MIT BMC for DNA sequencing; S. Vos for use of lab equipment and advice on protein purification; J. Gelles, L. Friedman, D. Price, and members of the G. - W.L. lab for discussions; R. Landick, P. Babitzke, K. J. Dierksheide, J. Xue, I. Kim, and E. Wirachman for comments on the manuscript. ChatGPT and Claude were used for copy editing.

## Author contributions

Conceptualization: RAB, GWL, EE, EOO

Methodology: RAB

Investigation: RAB, EE

Visualization: RAB, GWL

Funding acquisition: GWL, RAB, EOO

Writing – original draft: RAB, GWL, EE

Writing – review & editing: RAB, GWL, EE, EOO

## Data, code, and materials availability

Supplementary data tables; core scripts used for 3′-end sequencing analysis, decision tree model training and evaluation, and HUBfinder terminator prediction will be available upon publication.

## Supplementary Materials

Materials and Methods

Supplementary Text

Figs. S1 to S7

Tables S1 to S7

## Materials and Methods

### Strains

For purification of *B. subtilis* RNAP, a *rpoE rpoY* double knock-out strain containing a 6X-His C-terminal tag at the native *rpoC* locus in *Bacillus subtilis* subsp. subtilis str. 168 was generated (GLB638, Table S1). This strain was designed to remove the δ and ε accessory factors from our protein preparation, thereby mitigating the potential effects of heterogenous RNAP subunit composition on our experiments. *rpoE rpoY* double knock-out was generated by transforming gDNA from the BKK and BKE collection strains (BGSC, (*63*)), respectively, using *B. subtilis* natural competency (*64*). The resulting double knock-out strain was further transformed with pPolHis1 plasmid (BGSC, (*65*)) to add sequence encoding an in-frame 6X His-tag to the 3′-end of the *rpoC* gene.

### Template library generation

#### General template design

All *in vitro* transcription templates consist of a Pspank promoter (*66*) followed by a 29-bp sequence (A29) containing a single T base at +2 (relative the transcription start site, +1). The A29 sequence allowed the formation of halted elongation complexes during single-round *in vitro* transcription reactions (*22*). Immediately downstream of A29 was a 5-bp restart sequence (*22*) followed by a SalI restriction site. 90-bp variable terminator sequences were cloned downstream of the SalI site, followed 29 bps downstream by the invariant *trnB* terminator sequence.

#### Endogenous Terminators

Intrinsic terminator sequences were selected from *B. subtilis* and *E. coli* using end-enriched sequencing (Rend-seq) data from these organisms (as described in (*13*)). Briefly, secondary structures were predicted for sequences upstream of 3′-end peaks in genomic regions with sufficient coverage. Sequences with the following structure/sequence characteristics were chosen as putative intrinsic terminators: hairpin free energy < -7 kcal/mol, loop length between 3-10 nts, stem length between 4-17 bps, and a U-tract length of at least 2 U bases. After applying these thresholds, 1,413 and 598 intrinsic terminator sequences were chosen from *B. subtilis* and *E. coli*, respectively (Table S5). For each terminator, a sequence window spanning 60 bps upstream, including the position of the Rend-seq 3′-end peak, and 30 bps downstream of the peak were selected for the template library.

ssDNA oligos containing the A29 sequence, restart, SalI site, variable terminator sequence, and a downstream 30 nt homology arm were synthesized using phosphoramidite chemistry (Twist Biosciences). The oligo library was PCR amplified with primers oRAB372 and oRAB373 and Gibson cloned into a vector backbone between the Pspank promoter and *trnB* terminator sequences. Ligation product was cleaned and de-salted with a DNA Clean & Concentrator 5 column (Zymo Research) and transformed into 10-beta competent cells (NEB) via electroporation. Transformant colonies were grown overnight at 37°C on LB-agar plates with 100 μg/mL carbenicillin and scraped off into 10 mM MgSO_4_. Template library plasmids were extracted using ZymoPure Midiprep (Zymo Research).

#### Mutant Terminators

Our mutant intrinsic terminator template library spanned 5 categories: point mutations, -11/-10 position mutations, -2/-1 mutations, -11/-10 and -2/-1 combinations, and λtR2 mutations (described in Supplementary Information, Table S4 and S6). For point mutations within the predicted hairpin, versions with and without a complementary mutation on the opposite side of the hairpin were included. Likewise, for all -11/-10 position mutants, complementary mutations were made in the upstream half of the hairpin.

The mutant oligo library was synthesized and cloned as for endogenous terminators, but included a 10-nt unique barcode downstream of the SalI site. Barcodes were designed to have less than 6 T bases and be separated by a Hamming distance of at least 2 from all other barcodes in the library. Selection after transformation of the mutant library was done in 50 mL LB media with 100 μg/mL carbenicillin instead of on plates.

### *In vitro* transcription

*Template immobilization.* For all *in vitro* transcription reactions, template was bead immobilized to facilitate capture of released (i.e. terminated) RNAs. Linear DNA templates were generated using PCR of the plasmid library with a 5′-biotinylated forward primer (oRAB212, Table S2) and cleaned up with a DNA Clean & Concentrator 5 column (Zymo Research). Resulting product was attached to Dynabeads MyOne Streptavidin C1 magnetic beads (Invitrogen) using manufacturer’s instructions for RNA applications. At the final step, beads were resuspended in 1X TGA buffer (40 mM Tris Acetate pH 8.0, 100 mM potassium acetate, 8 mM magnesium acetate, and 4% glycerol) to a template concentration of 250 nM.

#### Holoenzyme formation

For *B. subtilis* RNAP, holoenzyme was formed by incubating 1 µM core RNAP with 3 µM σ_A_ in 1X TGA + 1 mM DTT at 37°C for 10 minutes.

#### Transcription Reaction

Halted elongation complexes (HECs) were generated by incubating 50 nM *B. subtills* or *E. coli* (NEB) holo RNAP with 50 nM bead-immobilized template in 1X TGA with 1 mM DTT; 5 µM ATP, CTP, and GTP; 100 µM ApU; and 1X Superasin RNase inhibitor (NEB) at 37°C for 15 minutes before placing on ice. For experiments with NusA and NusG, 1 µM NusA or NusG was incubated with the HECs for 5 minutes at 37°C prior to the resumption of elongation. For experiments with 2 µM NTPs, HEC reaction buffer was exchanged for 1X TGA with 1 mM DTT and 1X Superasin using magnetic separation before the addition of chase buffer. Elongation was restarted by the addition of chase buffer (1X TGA with 200 or 2 µM of each NTP and 20 µg/mL rifampicin) and incubation at 37°C for 15 minutes. Released RNAs were selected by taking the supernatant after magnetic separation of template-coated beads. The supernatant was combined 1:1 with 2X stop buffer (10 mM EDTA, 1% SDS, 40 mM Tris-HCl pH 8.0) to end all active transcription.

### Sequencing library preparation

3′-end RNA sequencing libraries were prepared based on the protocol from (*35*).

#### RNA preparation

Released *in vitro* transcribed RNAs in 1X stop buffer were extracted using acidic phenol:chloroform and ethanol precipitated. Precipitated RNA was resuspended in 10 mM tris pH 7.0.

#### Adaptor ligation

In T4 RNA ligase 2 reaction buffer (NEB) with fresh PEG 8000 (NEB), RNA was combined with 100 pmol of an adenylated adaptor (linker-I, Table S2) in at least a 3 pmol:100 pmol RNA:adaptor ratio. The ligation reaction with T4 RNA ligase 2, truncated K227Q (NEB) was allowed to proceed at 25°C for 5 hours. RNA was precipitated with isopropanol and gel purified using a 6% Urea-PAGE gel (Invitrogen). Ligated terminated and read-through (i.e. terminated at the downstream *trnB* terminator) products were gel extracted and kept together for all subsequent steps.

#### RT and Final PCRs

Ligated RNA was reverse transcribed using SuperScript III (ThermoFisher) with a primer complementary to the ligated adaptor and containing a 16 nucleotide-long unique molecular identifier (UMI) sequence (oJT442, Table S2). Following reverse transcription, RNA was hydrolyzed by incubating at 95°C with 100 mM NaOH for 5 minutes. Resulting cDNA products were size-selected with a 6% Urea-PAGE gel. Adaptors and indexes for sequencing were added to cDNA via two PCR steps (PCR 1 primers: oRAB374 and S1 indexing primer. PCR 2: S2 indexing primer and oRAB338, Table S2). cDNA was amplified for 8-11 cycles (4 cycles of PCR1 and additional PCR2 cycles) using Phusion DNA polymerase (NEB) and size-selected on an 8% TBE gel or by 2 rounds of magnetic bead clean up using a 1:1.5 DNA:bead ratio (PCRClean DX bead, Aline Biosciences). Resulting libraries were sequenced with 50-bp paired-end reads on Singular G4.

### 3′-end sequencing data processing

*Adaptor trimming and alignment.* UMIs were extracted from reads with UMI-tools (*67*) using parameters “--extract-method=string --bc-pattern=NNNNNNNNNNNNNNNN” followed by adaptor trimming with Cutadapt (*68*) using parameters “-g ATTGATGGTGCCTACAG”. Trimmed reads and their corresponding pairs were aligned to a fasta file of template sequences with Bowtie v1.2.3 using parameters “--no-unal”. SAM alignment files were converted to BAM files using Samtools (*69*) and UMI collapsed using UMICollapse (*70*) with “--remove-unpaired” and “--remove-chimeric” commands. Collapsed alignment files were converted to wig-type files of 3′-end counts using Bedtools (*71*) with parameters “-d -5 -strand -”.

#### Termination site identification

For each terminator in our template library, the termination site where the majority of termination occurred was determined by finding the position with the most 3′-end counts in a window spanning 9 bps upstream and downstream of the *in vivo* 3′-end (*13*). The same approach was used to determine terminator adjacent *in vivo* 3′-end peaks from *B. subtilis* WT and 4-exo strains (*28*).

#### Termination efficiency determination

Site-specific termination efficiencies (TE) were determined for positions in a window spanning 9 bps upstream and downstream of the *in vivo* 3′-end for each terminator. To determine site-specific TEs, the ratio of 3′-end counts at a site *j* (*S_j_*) to the sum of 3′-end counts at *j* and downstream (∑*^n^_i=j_ S_i_*) was calculated:

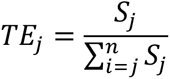

Total TE (*TE*_total_) for each terminator was calculated as:

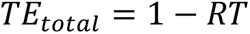

where RT is the total read-through of the terminator, calculated as the ratio of 3′-end counts in a 10-nt window around the *trnB* termination site to total 3′-end counts mapped to the corresponding terminator construct.

#### Contribution to total termination efficiency (*C*_TE_)

The contribution of a termination site to *TE_total_* was calculated as the fraction of terminated 3′-ends. To determine the fraction of terminated 3′-ends at a position, the ratio of 3′-end read counts at that site (*S_j_*) to the sum of 3′-end counts in a window spanning 9 bps upstream and downstream of the *in vivo* 3′-end (*T_sum_*) was calculated:

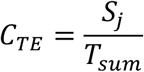

#### Sequence logo generation

Sequence logos for different TE tiers were generated for sequence windows (15 bps upstream and 10 bps downstream) relative to processed *in vivo* 3′-end peaks or *in vitro* termination sites. Tiers were structured using either total TE or site-specific TE. Sequences associated with TEs ≥ 0.90 were aggregated into fasta files and sequence logos were generated with WebLogo 3 (*72*). Background GC-content was set to 45.5% to account for the distribution of *B. subtilis* and *E. coli* terminators in our library and the composition of their respective genomes.

### Protein purification

*B. subtilis RNAP*. RNAP was purified out of *B. subtilis* strain GLB638. Cells were grown to OD_600_ 0.9-1.0 in LB supplemented with 1% glucose. Pelleted cells were resuspended in lysis buffer (50 mM Tris-HCl pH 7.9, 500 mM NaCl, 20 mM Imidazole, 1 mM DTT, 5% glycerol) and lysed using sonication. Lysate was passed over a HisTrap Ni-NTA column (Cytiva) and bound protein was washed with lysis buffer. RNAP was eluted from Ni-NTA onto a HiTrap heparin column (Cytiva) with elution buffer (50 mM Tris-HCl pH 7.9, 50 mM NaCl, 300 mM Imidazole, 1 mM DTT, 5% glycerol) and eluted from heparin using a continuous gradient from low salt buffer (50 mM Tris-HCl pH 7.9, 50 mM NaCl, 0.1 mM EDTA, 1 mM DTT, 5% glycerol) to high salt buffer (50 mM Tris-HCl pH 7.9, 1 M NaCl, 0.1 mM EDTA, 1 mM DTT, 5% glycerol). *B. subtilis* RNAP elutes from heparin in two peaks: a holoenzyme peak (⍺_2_ββ′ω +σ_A_) and a core peak (⍺_2_ββ′ω). The core peak was dialyzed into size exclusion buffer (20 mM Tris-HCl pH 7.9, 150 mM NaCl, 0.1 mM EDTA, 1 mM DTT, 5% glycerol) overnight. Core RNAP was further purified using size exclusion chromatography (Superose 6 10/300 GL, Cytiva) and dialyzed overnight into storage buffer (20 mM Tris-HCl pH 7.9, 250 mM NaCl, 1 mM MgCl_2_, 20 µM ZnCl_2_, 0.1 mM EDTA, 0.5 mM DTT, 20% glycerol) before concentrating and storing at -80°C for future use.

#### B. subtilis σ_A_

σ_A_ was purified as in (*73*). Briefly, *E. coli* Rosetta 2 cells expressing *B. subtilis* σ_A_ with a C-terminal intein-chitin tag were grown to OD_600_ 0.5 in LB media at 28°C, induced with 0.5 mM IPTG, and grown for 3.5 hrs at 25°C before harvesting cells. Cells were lysed by sonication in buffer A (25 mM Tris-HCl pH 8.0, 500 mM NaCl, 0.1 mM EDTA, 1 mM MgCl_2_, 10% glycerol) with 0.05% Triton X-100 and protease inhibitor tablet (Thermo). Clarified lysate was applied to chitin resin (NEB) for 1 hr with gentle agitation, drained, and washed with buffer A. Column was washed with buffer B (25 mM Tris-HCl pH 8.0, 100 mM KCl, 0.1 mM EDTA, 1 mM MgCl_2_, 10% glycerol) then incubated in buffer B with 50 mM DTT overnight to cleave the intein tag. Cleaved protein was eluted in buffer B, loaded onto a HiTrap Q HP column (Cytiva), and eluted via a gradient from 100 mM KCl to 500 mM KCl over 90 min. Peak fractions were dialyzed into storage buffer B (buffer B with 1 mM DTT and 20% glycerol) and stored at -80°C for later use.

#### B. subtilis NusA

*E. coli* Rosetta 2 cells expressing *B. subtilis* NusA with an N-terminal 10X-His tag were grown to OD_600_ 0.4-0.6 in LB media at 37°C and induced with 1 mM IPTG before shifting to 16°C overnight. Harvested cells were resuspended with lysis buffer (50 mM Tris-HCl pH 7.9, 300 mM NaCl, 20 mM Imidazole, 1 mM DTT, 5% glycerol) and lysed using sonication. Lysate was applied to a HisTrap Ni-NTA column (Cytiva), washed with lysis buffer, and eluted (elution buffer: 50 mM Tris-HCl pH 7.9, 50 mM NaCl, 300 mM imidazole, 5% glycerol) onto a heparin column before continuous gradient elution from low salt buffer (50 mM Tris-HCl pH 7.9, 50 mM NaCl, 1 mM DTT, 5% glycerol) to high-salt buffer (same as low salt, except 1 M NaCl). Eluted protein was dialyzed overnight into size exclusion buffer (20 mM Tris-HCl pH 7.9, 150 mM NaCl, 1 mM DTT, 0.1 mM EDTA, 5% glycerol) with TEV protease to remove His tag. Protein was flowed over Ni-NTA to remove cleaved His tag and TEV protease before a final size exclusion purification step (Superdex 75 10/300 GL, Cytiva). Protein was dialyzed overnight into storage buffer (20 mM Tris-HCl pH 7.9, 250 mM NaCl, 1 mM DTT, 0.1 mM EDTA, 20% glycerol), concentrated, and stored at -80°C for future use.

#### B. subtilis NusG

*E. coli* Rosetta 2 cells expressing *B. subtilis* NusG with an N-terminal 10X-His tag were grown to OD_600_ 0.4-0.6 in LB media at 37°C and induced with 1 mM IPTG before shifting to 16°C overnight. Harvested cells were resuspended with lysis buffer under denaturing conditions (50 mM Tris-HCl pH 7.9, 500 mM NaCl, 20 mM Imidazole, 1 mM DTT, 5% glycerol, 6 M guanidine HCl) and lysed using sonication. Lysate was applied to a HisTrap Ni-NTA column (Cytiva), washed with lysis buffer, and refolded on-column by a linear gradient from 6 M to 0 M guanidine HCl over 2 hours at 0.5 mL/min. flow rate. Refolded protein was eluted (elution buffer: 50 mM Tris-HCl pH 7.9, 500 mM NaCl, 300 mM imidazole, 5% glycerol) and dialyzed overnight into size exclusion buffer (20 mM, 500 mM NaCl, 1 mM DTT, 5% glycerol). A final purification step was conducted with size exclusion chromatography (Superdex 75 10/300 GL, Cytiva) before concentrating and mixing 1:1 with 2X storage buffer (20 mM Tris-HCl pH 7.9, 500 mM NaCl, 0.2 mM EDTA, 1 mM DTT, 40% glycerol). Protein was stored at - 80°C for future use.

### Decision tree modeling of termination efficiency

#### Hairpin folding

The best hairpin structure for each termination site was found by successively folding and scoring larger sequence windows relative to the termination site (starting at position -20 and proceeding downstream to -70). Positions -10 to -1 are constrained from folding to reflect their location in the DNA-RNA hybrid of RNAP. Scoring criteria incorporates minimum free energy (*mfe*), number of bulges (*bulges*), loop size (loop penalty, *penalty_loop_*), and stem length (*n_stem_*). Secondary structures and *mfe* were determined using the RNAfold package from ViennaRNA (*74*). Loop penalty for loops <4 nts was computed as:

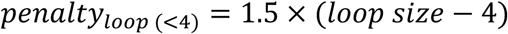

Or, for loops >4 nts:

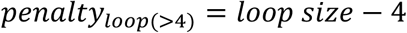

Late penalty (*penalty_late_*) was computed as:

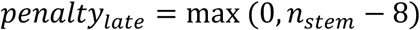

Score for each hairpin was computed as:

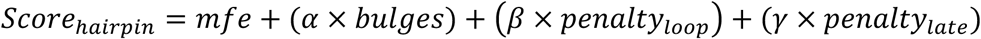

For feature extractions in this study the values of 1.0, 0.6, and 0.3 for *α, β*, and *γ*, respectively, were used to reflect the relative impact of bulges, loops, and stem length on terminator hairpin folding and function: An issue with using hairpin stability alone to choose the “best” hairpin at given site is that excessively long hairpins with discontinuous base pairing and/or large loops can have lower *mfe* values than shorter terminator hairpins with continuous pairing. These long hairpins are often unable to continue folding into the RNA-DNA hybrid, as would be expected for a true terminator hairpin. The values chosen for the hairpin scoring parameters above were selected by visual inspection to eliminate predicted hairpins that do not extend into the hybrid region (Hairpin Growth = 0, described below) for termination sites with TE>0.5. Additionally, when choosing the best scoring hairpin for each site, only single-stem folds were considered with a preference for folds where the edge of the hybrid (position -11) was base paired.

#### Feature extraction

For feature extraction, terminator sequences passing a threshold of 10^3.5^ total reads were used. From these terminators, the following features were extracted for sites starting at -9 relative to the *in vivo* 3′-end until the site where cumulative TE is >90% of total TE.

##### Hairpin features

- Hairpin ΔG – *mfe* value for hairpin folding.
- Hairpin Length – Number of base pairs in hairpin.
- Hairpin Growth – Number of continuous base pairs that can form into the hybrid sequence.
- Loop Sequence – Sequence of loop transformed using LabelEncoder() from sklearn.preprocessing.
- *U-tract features*.
- U-tract count – Number of U bases from -9 to -3.
- U-tract length – Longest continuous stretch of Us from -9 to -3.
- Hybrid ΔG – Free energy value for annealing of DNA-RNA hybrid calculated using values from (*75*).

##### Bubble sequence

- -11 to -1 – Unicode transformed nucleotide identity for -11 to -1 sequence.

##### Full-range sequence

- +41 to -19 – Unicode transformed nucleotide identity for +40 to -19 sequence.

Features for 24,220 sites from 2,006 different *B. subtilis* and *E. coli* terminators were ultimately used for model training and testing (Table S7). TE values <0.001 were set to 0.001 prior to training and evaluation.

#### Model Training

Select feature sets were used to train decision tree models using XGBRegressor from XGBoost. An 80/20 train/test split was applied to the data. TE values were logit transformed for training with an inverse logit function applied to the output for interpretability. The following parameters were used for training: n_estimators=200, max_depth=7, learning_rate=0.05, subsample=0.9, colsample_bytree=0.7, min_child_weight=5, and gamma=0.

#### Model evaluation

XGBoost models trained with different feature sets were used to make TE predictions for the designated test set of terminators. The correlation between predicted TE values and measured (i.e. true) TEs was evaluated by calculating R^2^. For a model trained on hairpin, U-tract, and bubble sequence, we further evaluated the deviation of predicted TEs from true TEs by calculating the log_10_ transformed ratio of predicted TE to true TE.

### Prediction of intrinsic terminators in bacterial genomes

#### HUBfinder terminator prediction

HUBfinder first collects stem, U-tract, and -11 to -1 sequence features corresponding to every site of a given genome. Next, our XGBoost model makes TE predictions for every site using the collected features and outputs the result as a .bedgraph file. From the site-specific predictions, the total TE for overlapping 4 nt windows is calculated. The position with the highest site-specific TE within a window that passes a threshold for total TE (default =0.4) is designated as the termination site for a putative terminator. Duplicate calls are filtered by enforcing a 10-nt gap between predicted sites.

#### HUBfinder evaluation

To assess whether bubble-edge sequence motifs (B) improve model performance across different bacterial genomes, we compared the genomic distribution of predictions and the consistency predictions with *in vivo* data between HUBfinder and a model that does not consider the bubble sequence (sequence at positions -11 to -1 relative to the termination site) in training the model (HUfinder).

Terminators were predicted in genomes from representative species of 15 different bacterial phyla (Table S3) using a 40% TE threshold. Prediction rates (predictions per kilobase) were compared between all non-coding regions downstream of annotated stop codons, extending until the next start codon or for 200 bp (after genes), and all intragenic regions (inside genes). Predictions after genes and inside genes were counted using bedtools map (*71*). To compare predictions from HUBfinder and HUfinder to *in vivo* 3′-ends mapped in *C. crescentus* (*13*), predictions were filtered to only include sites from regions with adequate coverage for *in vivo* peak-calling. Average read density in the neighboring region of each genomic position was calculated as the mean read counts across 50 nt windows upstream and downstream of the position. 2 nt gaps between a given position and each window were used. Only predictions at positions with an average density > .22 reads per million were used for downstream analyses. Bedtools closest (*71*) was used to find the distance between 3′-peaks and the closest prediction.

## Supplementary Text

### Mutant Library Description

All mutant terminator template sequences with a description of the mutation can be found in Table S5. The different general types of mutants are described below.

#### Point mutations

18 terminators (11 *B. subtilis*, 5 *E. coli*/*S. typhimurium*, and 2 phage terminators) with varying termination efficiencies (TEs) were used for mutational analysis.

- subtilis terminators: yvmA, serA, ftsZ, rplT, htpG, dhaS, treR, yhgE, proS, yojO, and ywpJ.
- coli/S. typhimurium terminators: rplQ, accD, fabB, yidF, and his
- terminators: λtR2 and t500

For each terminator, the site with the maximum TE in our endogenous terminator dataset or literature (λtR2: (*21*), t500: (*5*), *his*: (*59*, *76–78*)) was used to define a window extending from 11 bps upstream of the 5′ edge of the hairpin to 15 bps downstream the termination site. Each position within the window was substituted with the 3 alternative bases. The window was widened for terminators with multiple sites of termination.

#### -11/-10 mutations

For three *B. subtilis* (*dhaS, yvmA,* and *rplT*) and one phage terminator (λtR2), the -11/-10 position relative to the main termination site was substituted with every possible dinucleotide sequence (16 for each terminator).

#### -2/-1 mutations

Using the same terminators as above, the -2/-1 position relative to the main termination site was substituted with every possible dinucleotide sequence (16 for each terminator).

#### -11/-10 and -2/-1 combinations

For *dhaS, yvmA,* and *rplT* we additionally generated all possible combinations of -11/-10 and -2/-1 dinucleotides (256 for each terminator).

#### λtR2 mutations

In addition to the mutations described above, we also replicated 9 previously published λtR2 mutations (tR2-5, 6, 7, 11, 12, 13, 14, 16, and 17 (*36*), Table S4, Fig. S1G). The ratio of site-specific TEs between λtR2’s main termination sites—U7 and U8—are consistent between our sequencing-based measurements and the original gel-based measurements with these mutants. Results for tR2-5 and -7 are not reported because these mutations eliminated termination, as expected.

λtR2 has a GC dinucleotide at the upstream edge of the transcription bubble and an AU dinucleotide at the downstream edge. However, they are not in phase to form the ideal GC(N_7_)RU motif because they are separated by six intervening bases instead of seven. The two main termination sites—U7 and U8—for λtR2 position the RU at -2/-1 or the GC at -11/-10, respectively. We wondered whether insertions that properly phase these dinucleotides to form the GC(N_7_)RU motif would make λtR2 stronger. To test this, we designed mutants with every possible base insertion for a position upstream of the GC (will not change phase, but increases hairpin length) and for every position -10 to -3 in the hybrid region (will change phase). Insertions in the hairpin were complemented on the upstream half of the hairpin. Insertion of an extra U in the U-tract to form a near optimal GC(N_7_)RU sequence increased termination efficiency by ∼50% compared to WT (Fig. S4C). Insertions after GC, but before the start of the U-tract were generally unfavorable. Non-U insertions in the beginning of the U-tract also resulted in TEs weaker or similar to the WT sequence, indicating that discontinuities at certain positions in the U-tract are detrimental to termination. Accordingly, non-U insertions near the - 2A resulted in TEs similar to the U insertion.

---

**Fig. S1.**
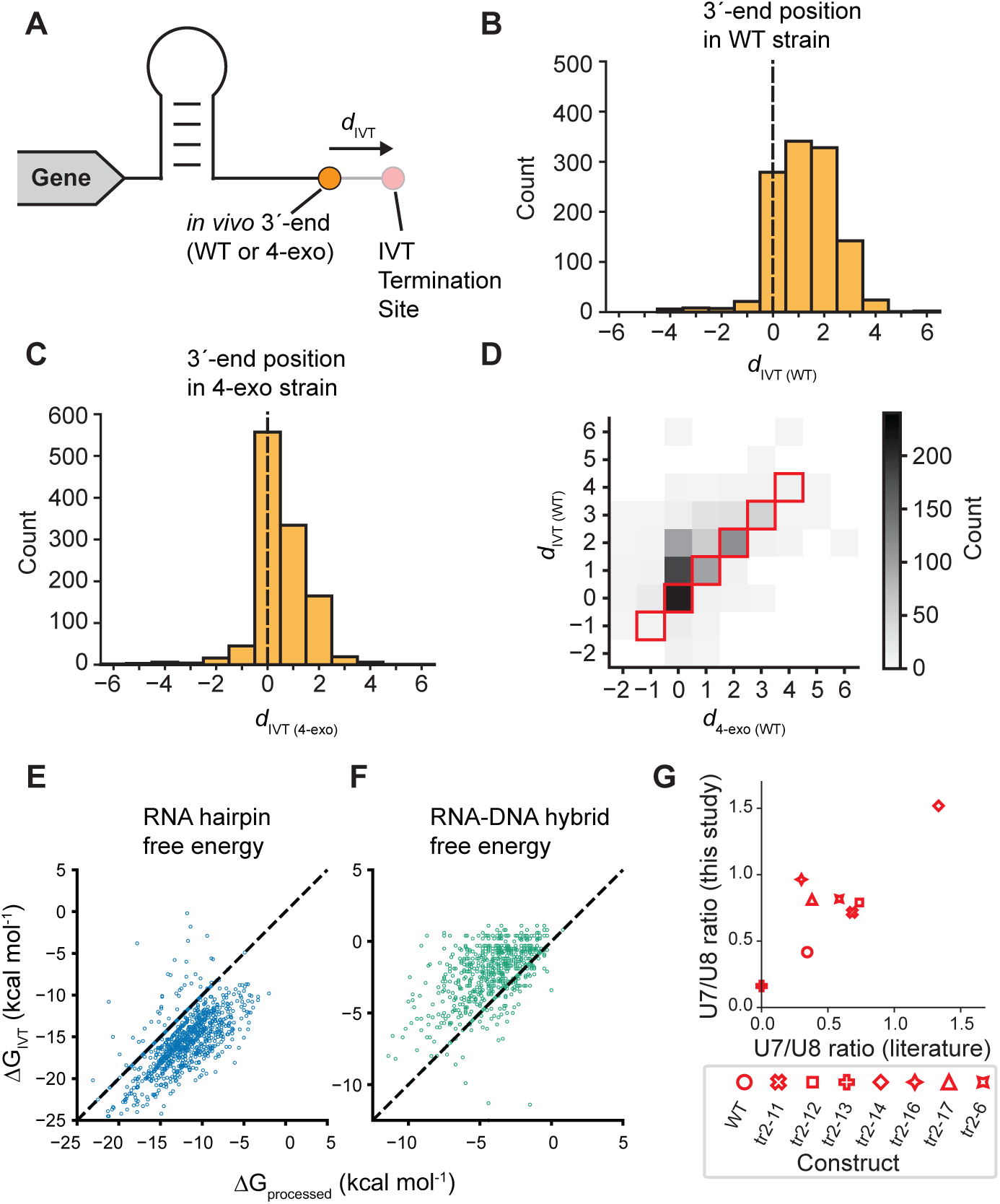
High-throughput measurement of *in vitro* termination sites. (**A**) *In vivo,* the positions of termination sites (red) and measured 3′-ends (orange) can be separated by some nucleotide distance (*d*_IVT_) because of processing by 3′-5′ exoribonucleases. (**B**) Histogram of nucleotide distances between *in vitro* termination sites and *in vivo* 3′-ends across intrinsic terminators in a WT strain of *B. subtilis* (*d*_IVT (WT)_). (**C**) Histogram of nucleotide distances between *in vitro* termination sites and *in vivo* 3′-ends across intrinsic terminators in a *B. subtilis* strain without 4 major exoribonucleases (4-exo) across intrinsic terminators (*d*_IVT (4-exo)_). (**D**) 2D histogram comparing the nucleotide distances between *in vitro* termination sites and WT *in vivo* 3′-ends (*d*_IVT (WT)_) to the nucleotide distances between 4-exo *in vivo* 3′-ends and WT *in vivo* 3′-ends WT (*d*_4-exo (WT)_). Counts on diagonal (red boxes) are terminators where *d*_IVT (WT)_ = *d*_4-exo (WT)_. (**E**-**F**) Hairpin (**E**, blue) and RNA-DNA hybrid (RNA-DNA duplex from position -10 to - 1) (**F**, green) stabilities measured relative to *in vitro* termination sites (ΔG_IVT_) plotted against the same features measured relative to *in vivo* 3′-ends (ΔG_processed_) for *B. subtilis* intrinsic terminators. (**G**) Comparison of termination measurements at U7 and U8 sites across λtR2 mutants between 3′-end sequencing (this study) and gel-based approaches (literature, (***24***)). The ratio of U7 and U8 TEs measured in this study and as in (***24***) are plotted against each other for different mutations. (denoted by unique symbols). Legend uses mutant names from (***24***) (as described in Table S4).

**Fig. S2.**
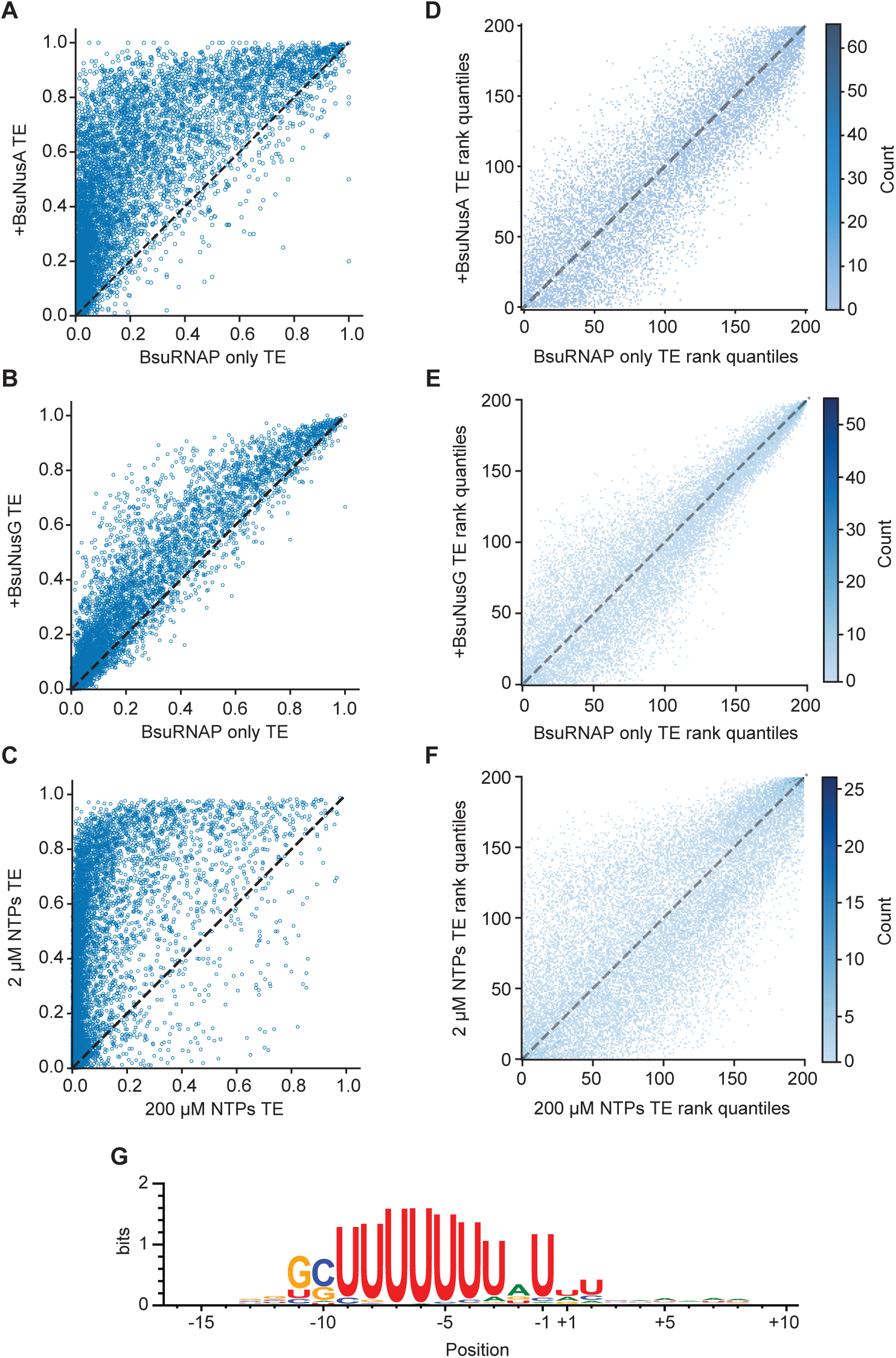
Termination efficiency with different transcription factors and RNAPs. (**A**) Scatter plot of site-specific TEs from transcription reactions with (+BsuNusA) and without (BsuRNAP only) NusA. (**B**) Scatter plot of site-specific TEs from BsuRNAP only transcription reactions with 2 µM and 200 µM NTPs. (**C**) Scatter plot of site-specific TEs from transcription reactions with (+BsuNusA) and without (BsuRNAP only) NusG. (**D**) 2D histograms comparing rank quantiles of site-specific TEs from transcription reactions with NusA (**D**, +BsuNusA), NusG (**E**, +BsuNusG), and 2 µM NTPs (**F**) to reactions with BsuRNAP only and 200 µM NTPs. (**G**) Sequence logo generated relative to termination sites (-1) with TE >= .90 as measured with *in vitro* 3′-end sequencing of an *E. coli* RNAP transcription reaction.

**Fig. S3.**
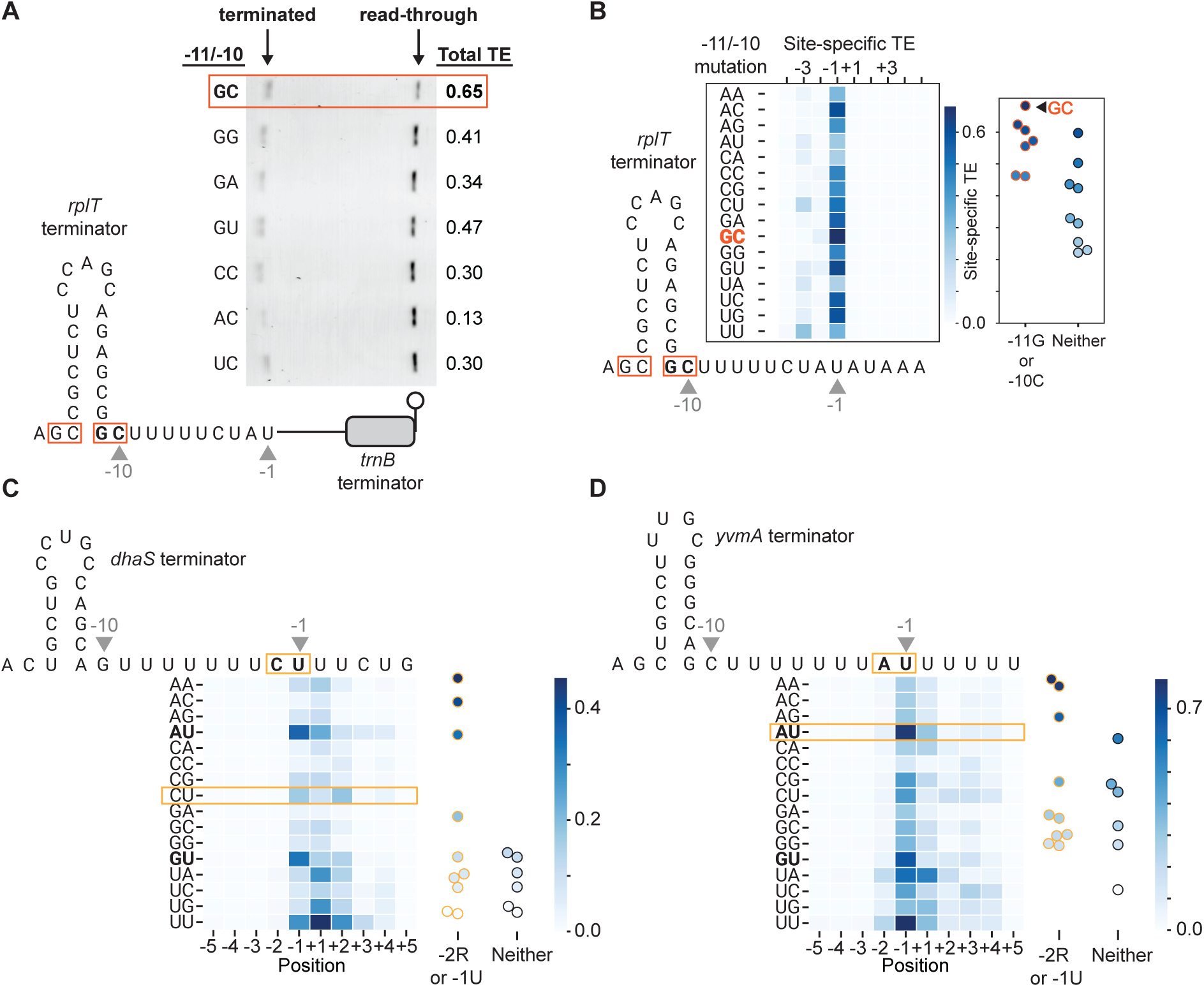
Additional -11/-10 and -2/-1 mutagenesis results. (**A**) Impact of mutagenesis on the -11/-10 position of the *B. subtilis rplT* terminator measured using a gel-based approach. Diagram indicates the positions mutagenized with red boxes. The nucleotide identity at -10/-11 for the template sequence associated with each lane is shown on the left side of the gel image. On the right side, the total TE (ratio of terminated signal to the sum of terminated and read-through signals where the signal is the band intensity normalized by length, Methods) is shown for each mutant. (**B**-**D**) -11/-10 mutagenesis (*rplT*, **B**) and -2/-1 mutagenesis of terminators lacking RU (*dhaS*, **C**) or with RU (*yvmA*, **D**) measured using 3′-end sequencing. WT residues are boxed on heatmaps, whereas consensus residues are bolded. Heatmaps display site-specific TEs in a 9-nt window around the -1 site for each mutant. Right inset: dot plots of site-specific TEs at -1 position for mutants with or without any motif matching sequences.

**Fig. S4.**
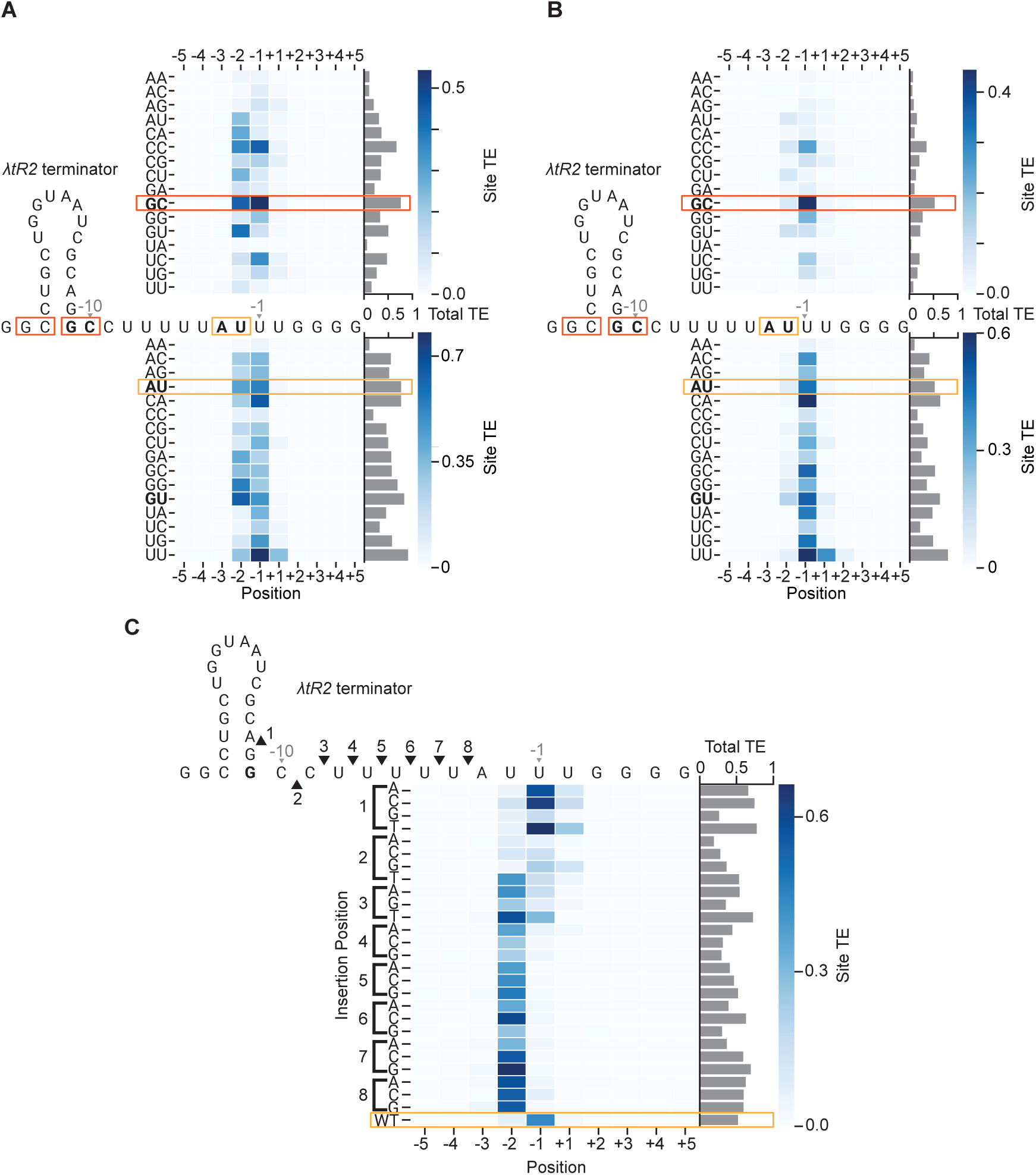
Additional λtR2 mutagenesis results. (**A**-**B**) Effects of mutagenesis to the -11/-10 (top) and -2/-1(bottom) positions of the λtR2 terminator transcribed by *E. coli* (**A**) and *B. subtilis* (**B**) RNAPs. Positions of mutagenesis are boxed (-11/-10, red and -2/-1, orange) on terminator sequence diagrams. WT residues are boxed on heatmaps, whereas consensus residues are bolded. Heatmaps display site-specific TEs in a 9-nt window around the -1 site for each mutant. Right inset: bar graphs of total TEs for each terminator sequence. (**E**) Effect of insertion mutations on the λtR2 terminator transcribed by *B. subtilis* RNAP. Nucleotide positions are relative to WT termination position (-1). Insertion locations are numbered 1 – 8 with their positions indicated on the sequence diagram with black triangles. Each row in the heatmap corresponds to a different nucleotide inserted at each indicated location. Right inset: bar graphs of total TEs for each terminator sequence.

**Fig. S5.**
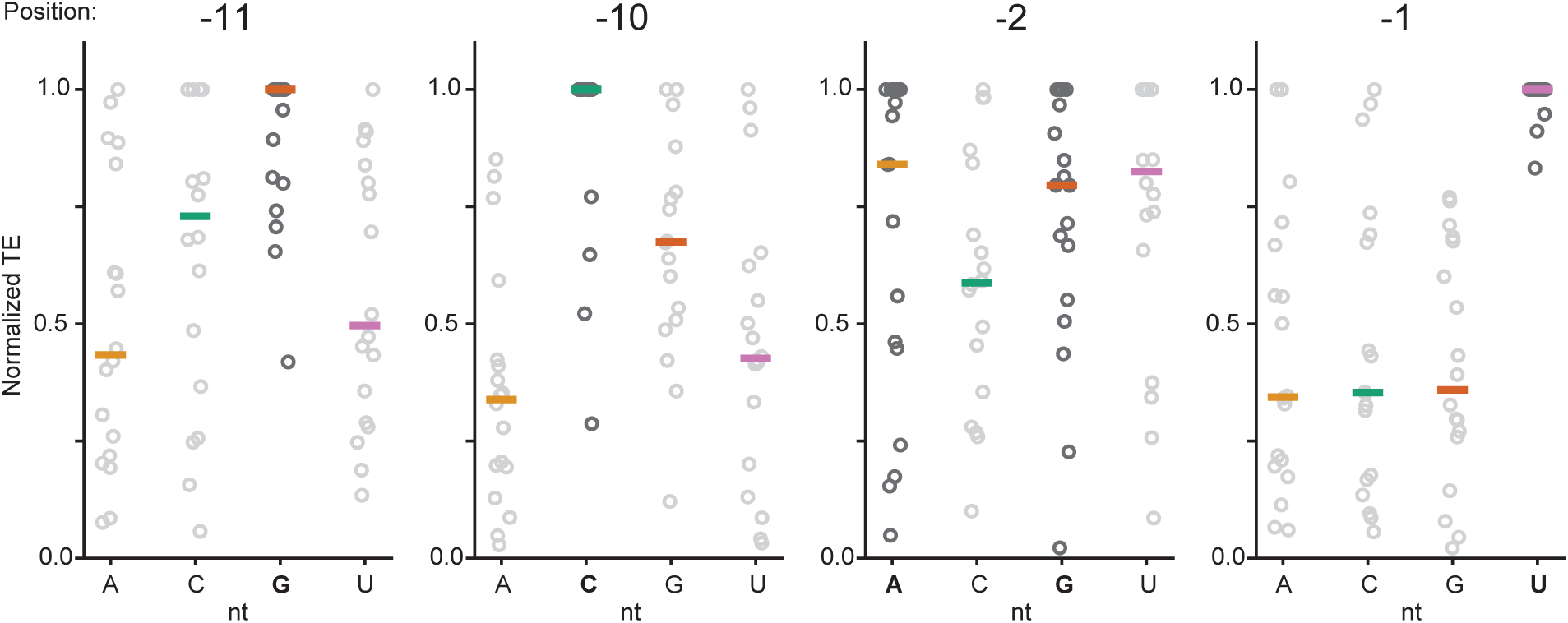
Mutagenesis targeting the bubble-edge sequences across 18 terminators. Effects of point mutations at template positions -11, -10, -2, and -1 (where -1 is the point of termination) for 18 *B. subtilis*, *E. coli,* and phage terminators. The -11 and -10 mutations are paired with compensatory mutations in the stem. Normalized site-specific TEs are plotted for variants with each nucleotide identity at the indicated position. For each terminator, TE is normalized to the max TE value at that position across the four possible nucleotide identities. Colored horizontal bars correspond to the median normalized TE for each nucleotide. Points and labels for nucleotides matching the consensus bubble-edge motif are bolded.

**Fig. S6.**
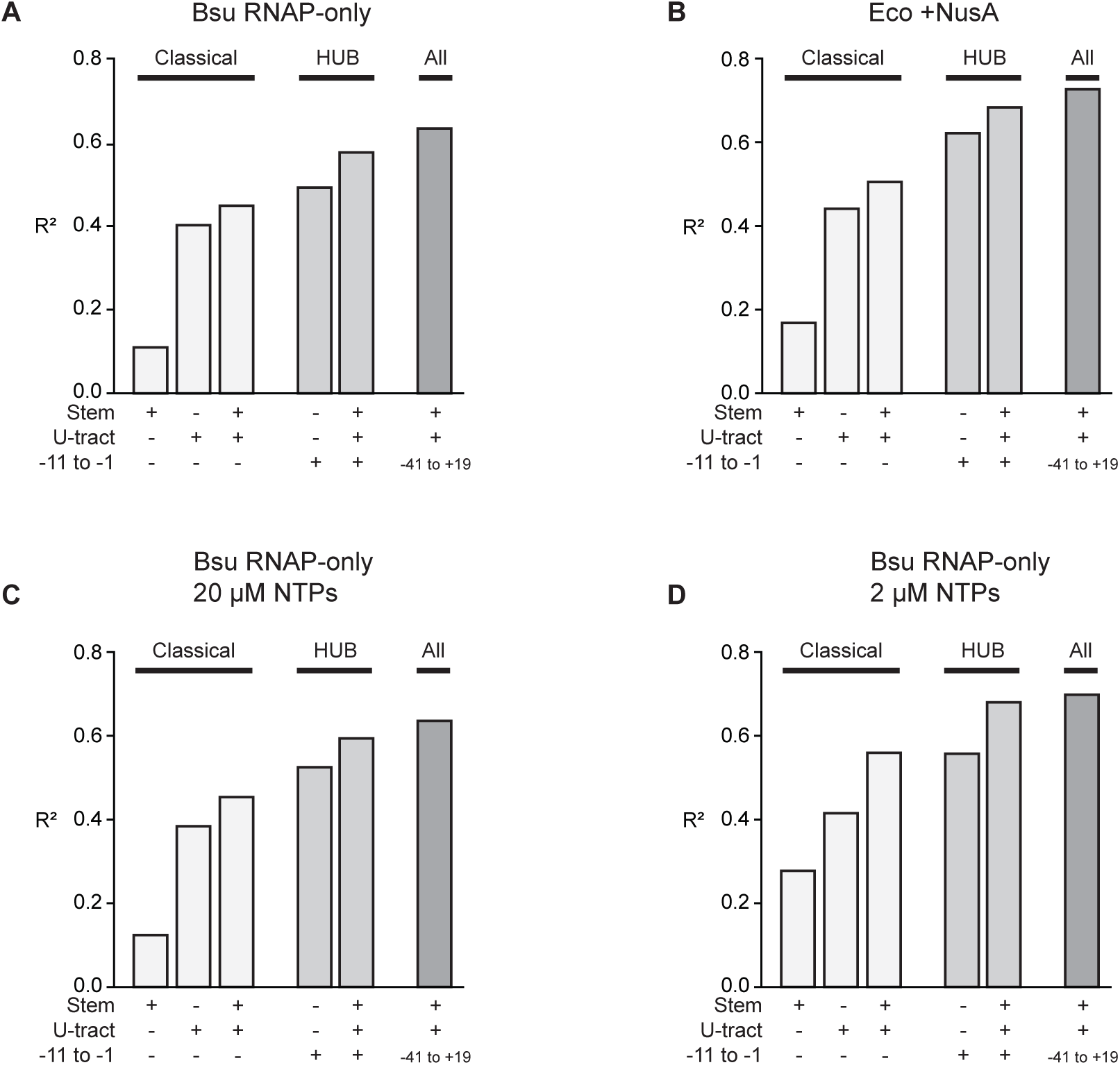
TE prediction model performance after training with different datasets. (**A**-**D**) Models that incorporate various HUB features for predicting TE trained using data from transcription reactions with *B. subtilis* (Bsu) RNAP only (**A**, with 200 µM NTPs), *E. coli* RNAP +NusA (**B**, with 200 µM NTPs), Bsu RNAP with 20 µM NTPs (**C**), and Bsu RNAP with 2 µM NTPs (**D**). Bar graphs plot the R^2^ values for statistical models trained with different feature sets. Classical feature set includes Hairpin and U-tract (Methods). The HUB feature set further includes Bubble sequence information (sequence of -11 to -1 region), whereas “All” features include sequence from an extended -41 to +19 region.

**Fig. S7.**
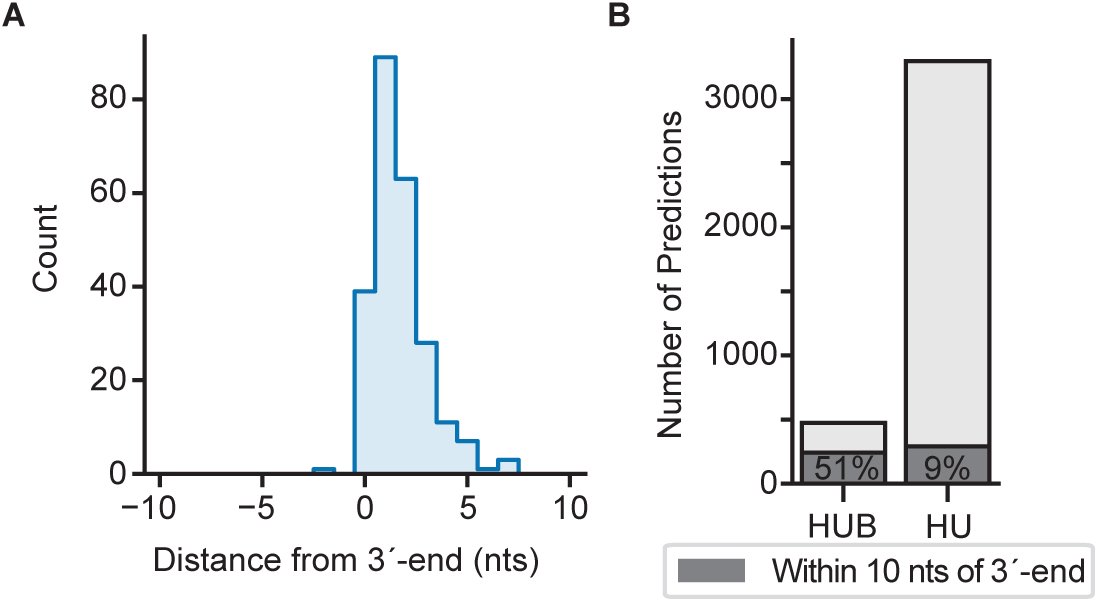
Distribution of HUBfinder and HUfinder predictions relative to *C. crescentus in vivo* 3′-ends. (**A**) Histogram of distances between HUBfinder predictions and 3′-end mapped in *C. crescentus* (*6*). Only predictions within genomic regions passing peak calling thresholds for Rend-seq analysis (average reads > .22 reads per million in 50 bp windows upstream and downstream of a given position, Methods) are considered. (**B**) Numbers of total HUBfinder and HUfinder predictions. Stacked bar graphs of total predictions from HUBfinder and HUfinder with dark grey area indicating predictions within 10 nts of a 3′-end. Percentage indicates the fraction of predictions that are within 10 nts of a *C. crescentus* 3′-end peak.

